# Sex-specific Flexibility in Breeding and Helping Strategies in the Sociable Weaver (*Philetairus socius*)

**DOI:** 10.64898/2026.01.28.702364

**Authors:** Jorge García-Campa, Liliana R. Silva, André C. Ferreira, Nicolas J. Silva, Franck Theron, Claire Doutrelant, Rita Covas

## Abstract

Helping-at-the-nest is often viewed as a precursor to reproduction, but switching between breeder and helper roles has been documented in some species. Such flexibility should depend on the opportunities and benefits of helping, which could differ between sexes due to dispersal strategies and social structure. However, whether breeding-helping flexibility is widespread and sex-specific remains unknown. Here, we investigated sex-specific strategies in breeding-helping flexibility using a 10-year dataset (1955 individuals) on sociable weavers (*Philetairus socius*), a colonial cooperative breeder in which males are typically philopatric whereas females disperse. Both sexes helped for several years, with males helping more frequently than females and for twice as long (0-13 vs 0-10 years). Around 40% of non-dispersing birds never became breeders and 50% of individuals bred without helping first. Both sexes switched roles between- and within-seasons but males were more likely to alternate (respectively four and six times more than females and up 15 switches across seven years). Our study reveals important flexibility and sex differences in breeder–helper roles, consistent with sex-biased dispersal, spatial breeding proximity and possible indirect fitness benefits. These factors could play a role promoting the evolution of helping across life-stages, not only pre-reproduction, but also reproductive and post-reproductive individuals.

## Introduction

Reproductive strategies can be fixed— e.g., annual breeding in many temperate vertebrates— or flexible, with individuals breeding multiple times, or skipping certain years [1-2]. Such flexibility across environmental and social contexts [3] allows them to optimize reproductive success by adjusting their behaviour or social roles over time [4-6]. Life-history traits and environmental variation are important drivers of this flexibility. For example, long-lived species often exhibit more conserved breeding strategies, or species exposed to fluctuating environments tend to exhibit higher plasticity [6-8].

In cooperative breeders, where individuals other than parents contribute to raising young [9], helping is usually considered a precursor to reproduction, in a relatively fixed path [10-11 but see [12]. However, cases of flexibility in changing from breeding to helping have been documented, with some long-term studies revealing that helping can arise after failed breeding attempts. For example, in long-tailed tits (*Aegithalos caudatus*), both male and female helpers are failed breeders that redirect their investment to raise nestlings belonging to other pairs [13] and in Iberian magpies (*Cyanopica cooki*), both sexes can help after unsuccessful breeding attempts, but only males help before reproducing [14]. Additionally, in superb starlings (*Lamprotornis superbus*), most individuals (73%) swap between breeder and helper roles bidirectionally throughout their life [15]. Together, these studies reveal a gradient of flexibility that remains poorly understood.

Flexibility in breeding and helping strategies is expected to differ between sexes when dispersal patterns, reproductive opportunities and benefits of helping diverge [16–19]. The philopatric sex has more opportunities to help kin or social partners and a higher likelihood of obtaining a breeding position within its natal group [19–22]. For example, in white-browed sparrow weavers (*Plocepasser mahali*), higher female helping is thought to reflect female philopatry and associated indirect fitness benefits [23]. Such sex-dispersal patterns are therefore expected to generate sex differences in flexibility, particularly in species reproducing in spatial proximity, where helping opportunities persist even after independent reproduction [24]. However, as with many traits that differ between sexes [25-26], breeding–helping flexibility has rarely been examined from a sex-specific perspective.

In this study, we use a 10-year dataset to investigate sex-specific flexibility in helping and breeding behaviour in the sociable weaver *Philetairus socius*, a colonial cooperative breeder with female-biased dispersal, resulting in genetic structuring among males but not females within colonies [27-28]. We describe individuals according to their reproductive categories (e.g., breeder, helper, non-breeder-non-helper) over the study period, and then examine sexual differences in i) helper dynamics (e.g., number of helping events, number of seasons helping, age at helping events, etc), and ii) breeder-helper dynamics (i.e., transitions between helping and breeding statuses). We simulate the breeding and helping trajectories to compare observed strategies with random expectations, under a null model assuming no sex differences. This comparison allows us to test whether the observed patterns reflect sex-specific reproductive strategies rather than chance. Moreover, because most females disperse to breed, we additionally identified a subset of resident females (i.e., those born and subsequently breeding in our study population; 25.7% of all females), to reconstruct complete reproductive trajectories and assess cooperative investment. We expected that females would concentrate their helping efforts toward their kin in their natal colony before dispersing, while males could more flexibly alternate between helper and breeder roles, given their opportunities to help kin or other familiar birds over their lifetime (i.e., before, during, or after reproducing).

## Material and methods

### Study area, model species and data collection

We monitored 27 sociable weaver colonies (10–130 individuals each) from 2014 to 2023 at Benfontein Nature Reserve, Northern Cape, South Africa (28°52’00’’S 24°51’00’’E).

Sociable weavers are endemic to the semi-arid savannahs of the Kalahari regions of Southern Africa and live in large communal nests used for both breeding and roosting throughout the year [29]. In our population, the breeding season is highly variable in timing and duration (4-11 months) [30]. These long breeding seasons allow some females to lay multiple clutches per season (mean clutch size = 3.17 ± 0.81; n = 4407; [31]), providing repeated opportunities to help and breed. Around 30-80% of nests are attended by groups composed of the parents plus 1-9 helpers, with helper numbers being influenced by previous year reproduction [32-34].

Before each breeding season (late August-early September), all individuals were captured, colour-ringed and blood sampled to genetically determine sex, parentage and relatedness. Nests were monitored every three days throughout breeding. Nestlings were individually marked and blood-sampled on day 9 for genetic analyses. Active nests were video-recorded during incubation (2-4h) and nestling period (usually for a total of 4-8h when the nestlings are ca. 9 and 17 days old). Breeding and helping roles were assigned using combined genetic and video data (following [30, 35]; see also *Determination of Reproductive Strategies* in SM).

### Classification of reproductive strategies

Individuals were classified into seven reproductive status categories: i) breeders (i.e., individuals observed breeding at least once but never helping) ii) helpers (i.e., individuals observed helping at least once but never breeding), iii) non-breeder-non-helper (i.e., individuals captured but never seen helping nor breeding); and iv) both helper and breeder (recorded both breeding and helping— whether or not in the same season). To reflect flexibility, we further subdivided the last category into four mutually exclusive categories reflecting temporal dynamics: a) helper-breeder, b) breeder-helper, c) helper-breeder-helper, and d) breeder-helper-breeder. These categories capture the sequence of role switches between helping and breeding observed in individuals throughout the study period. Individuals were considered potential helpers only from 150 days old, ensuring nutritional independence and meaningful contributions to nestling care (coinciding with adult plumage acquisition).

To confirm individual presence without helping or breeding, we combined video observations with annual capture data (see above). Individuals were assumed present if captured in the previous and recaptured in the subsequent annual capture sessions.

### Individual flexibility in cooperative and reproductive strategies

#### Helping dynamics

For each individual, we recorded i) the number of helping events, i.e., breeding attempts where it provided care to a nest other than its own, as well as ii) the number of breeding seasons in which they were observed helping. We also calculated the age at each helping event, allowing us to calculate iii) the age when individuals first helped and iv) the average age during helping events (following [36]; criteria for age calculation; see ESM).

#### Transition between helping and breeding statuses

We assessed whether i) an individual could help while simultaneously breeding. For this, we considered helping events as occurring in temporal proximity to breeding when the date of egg laying in the individual’s own nest was within 15 days before or after the helping event (i.e., potentially co-occurring with incubation or nestling provisioning). ii) We quantified the total number of transitions between helper and breeder statuses across the study period, as well iii) as the number of breeding seasons in which these changes occurred.

### Sex differences in Reproductive Strategies (Simulated vs. Observed)

To investigate whether males and females differ in their reproductive strategies from random expectations, we ran 10000 randomization simulations under a null model in which individual sex was permuted while keeping the total number of males and females constant. This generated expected distributions of reproductive strategies under the assumption of no sex-specific patterns. For each reproductive strategy, we report observed and mean of simulated percentages, standard deviations, and 95% confidence intervals.

## Results

We analysed 3759 reproductive events, comprising 3581 and 3408 breeder and 1652 and 629 helper records for males and females, respectively (individuals could contribute to several events; see ESM). In total, 958 males and 997 females (283 residents) were included in the analyses (Table S1).

### Helping dynamics

Using our full dataset (with all males and all females ever captured), we found that males were more likely to act as helpers than females in our population: 65.9% of all males helped at least once, while only 40.3% of females did (Fig. 1; Table S1). Simulations assuming that helping behaviour was randomly distributed across individuals predicted that 52.9% of individuals of both sexes would help at least once, so males help more than the expected by chance, and females less. When focusing on the data set containing only the resident females (those that were born and became breeders in our colonies, and hence for which we had the full life-history; n = 283) versus the immigrant females (n = 714), we found that 56.2% of resident females helped at least once in comparison to only 34% immigrant females (those that move to our colony to reproduce), so values for resident females were closer to males.

**Figure 1.**
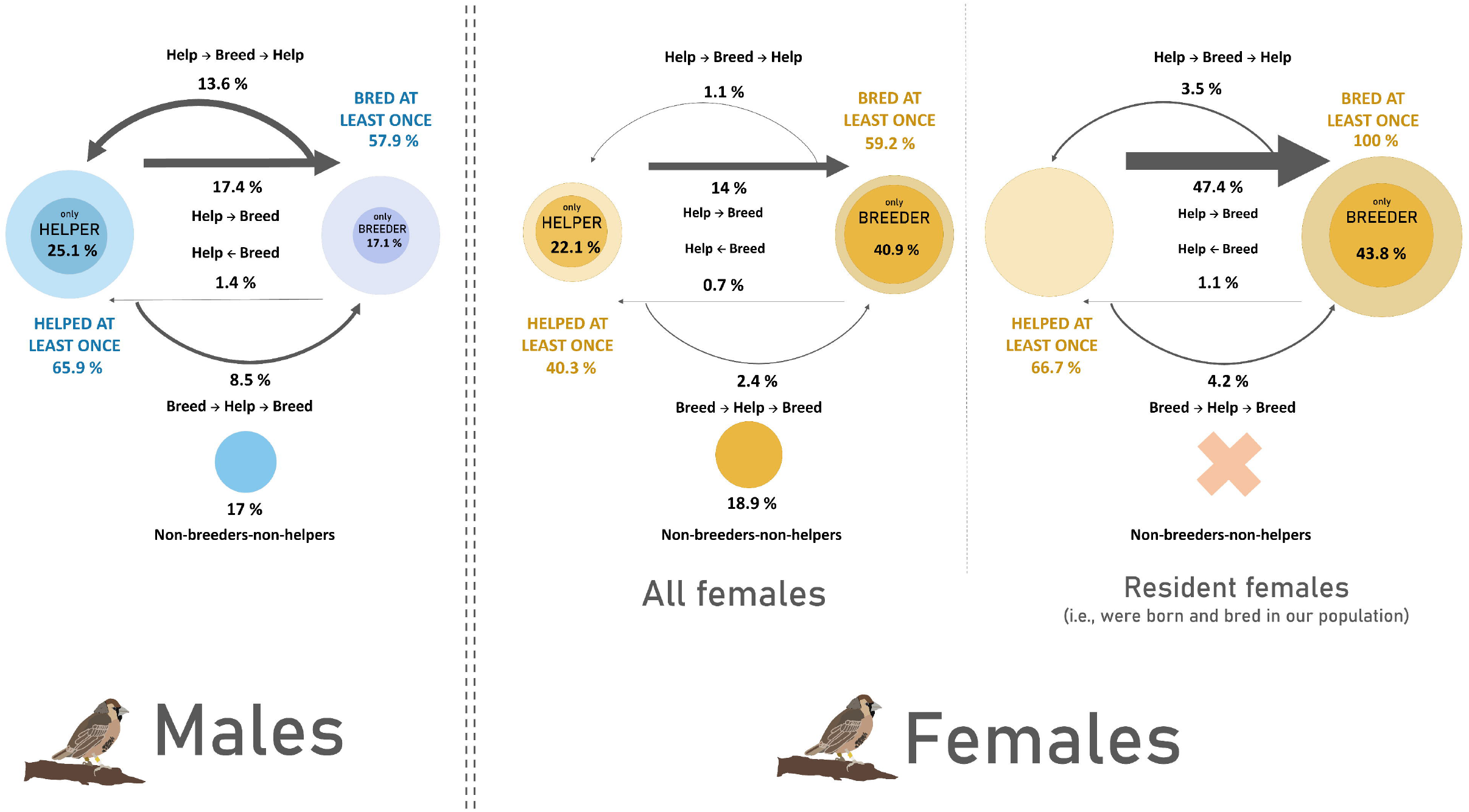
Percentages of reproductive categories observed from 2014 to 2023 for (i) males (n= 958), (ii) all females (immigrant + resident; n= 997), and (iii) resident females only (i.e., born and later breeding within the study population; n= 283). Circle sizes show the proportion of individuals recorded helping at least once, breeding at least once, or never observed helping or breeding. Inner circles represent exclusive categories (only helpers or only breeders, and arrow thickness indicate the percentage of individuals switching between roles (i.e., breeder→helper, helper→breeder, breeder→helper→breeder, or helper→breeder→helper). For resident females (born and breeding in the population), the non-breeder–non-helper and helper-only categories are not applicable, as indicated by the red cross and the absence of an inner circle, respectively. Colours denote sex (blue = males, orange = females).

Females concentrated their helping effort earlier in life and across fewer events than males. Using the full dataset, the age of first helping event was significantly higher for males (mean ± SD: 632 ± 579 days) than for females (408 ± 437 days; Welch’s t-test: t_(1003.4)_ = -4.10, p < 0.0001), and males helped in more than twice as many events as females (males: 2.08 ± 2.03; females: 0.77 ± 0.99; Mann-Whitney U test, W = 184294, *P* < 0.0001: Fig. 2a), and across more than twice as many breeding seasons (males: 1.28 ± 0.99; females: 0.56 ± 0.63; W = 186174, *P* < 0.0001; Fig. 2b). The most active male helper was observed across four breeding seasons (14 events), while the most active female was resident in our population and helped during three seasons (6 events). When using the dataset consisting only of the resident females, we found similar helping profiles to the overall female groups (mean helping events: 0.97 ±⍰1.11; mean seasons: 0.67⍰±⍰0.68).

**Figure 2.**
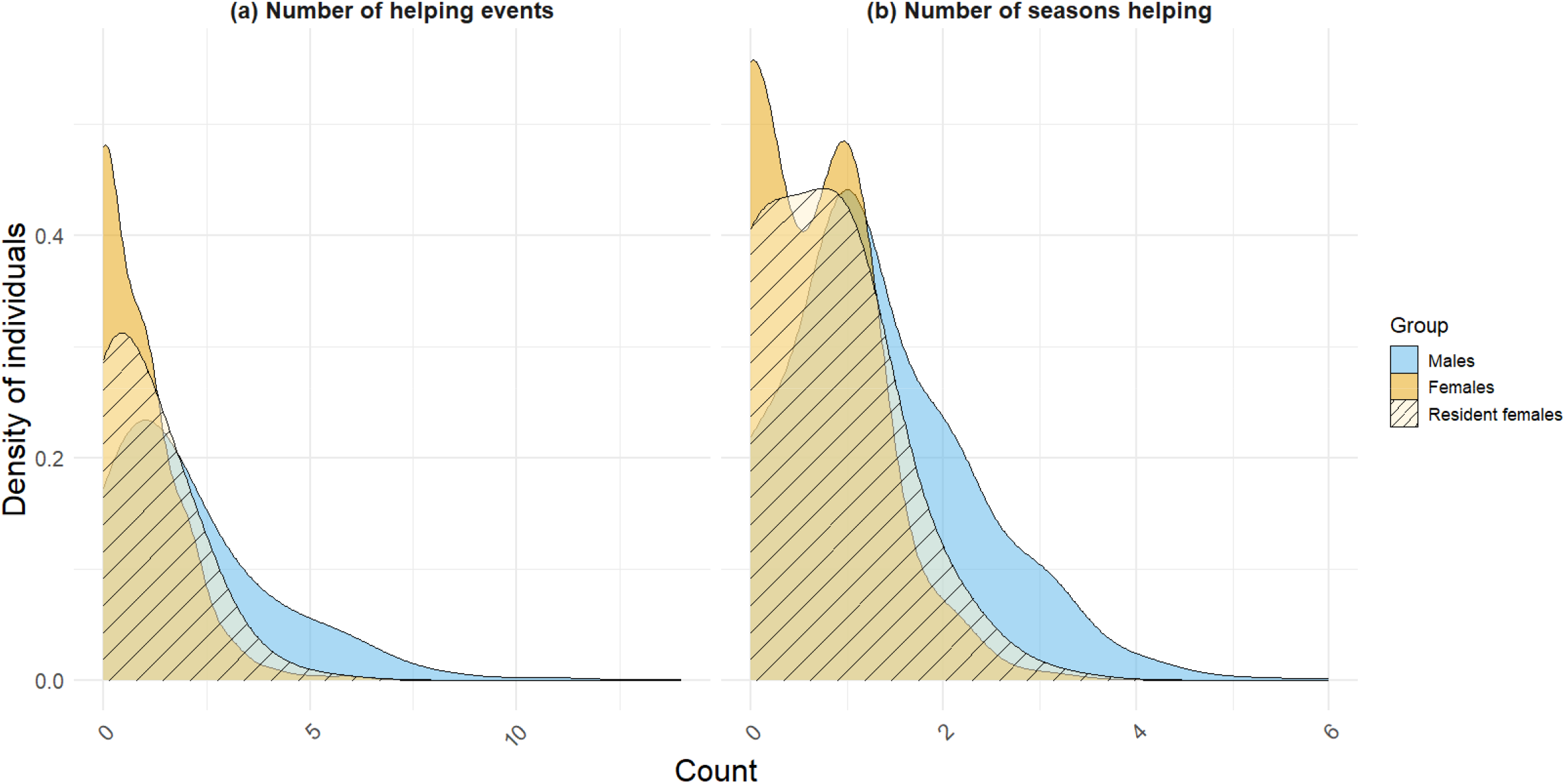
Number of helping events (a) and breeding seasons (b) in which individuals were observed helping. Data included all the individuals detected helping at least once. Densities are represented for all males (blue), all females (yellow), and resident females (pale yellow).

Males distributed their helping efforts more broadly across life. The average age during helping events was almost twice as high in males (748 ± 609 days) as in females (471 ±450 days; Welch’s t-test: t _(1527.1)_ = –11.88, p < 0.0001). Some individuals were seen helping over a decade, with records up to ∼13 years in males and ∼10 years in females (4686 and 3603 days, respectively). Among resident breeding females, the mean age during helping events was 629 ± 450 days (range = 151–2870 days, ∼8 years).

Simultaneous helping and breeding were rare, particularly among females. Only 1% of female helping events (11 of 1074) occurred during an active breeding attempt, compared to 4.7% in males (96 of 2057).

### Transitions between breeding and helping behaviour

A similar proportion of males and females reproduced at least once (57.9% and 59.2%, respectively; note that for the purpose of this study resident females were the ones that reproduced in the study colonies; Fig. 1). The proportion of individuals that remained non-breeder-non-helpers was also comparable between sexes: 17% in males, 18.8% in females (simulated ≈17.9%; Fig. 3). By contrasts, males were less frequently classified as ‘only breeders’ (17.1%) than females (40.9% for all females, 43.8% for the resident females) (Fig. 1 and 3; Table S1), contrasting with our simulations of 29.3% expected for both sexes (Fig. 3: Table S1).

**Figure 3.**
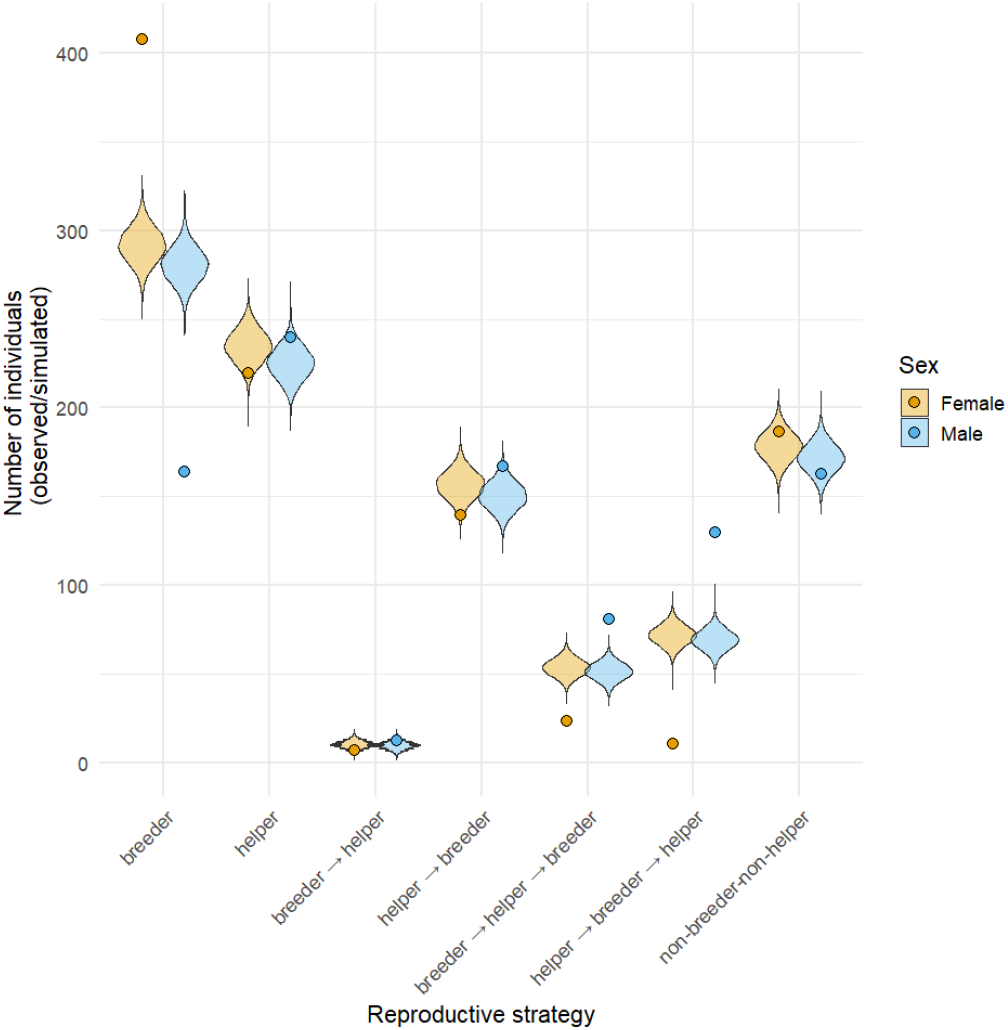
Observed and simulated numbers of individuals for each reproductive strategy in sociable weavers. Violin plots show expected counts from 10,000 simulations under the null model. Points indicate the observed values. Colours designate sex (blue = all males, orange = all females).

Helping was not a prerequisite for becoming a breeder in either sex. Among males that reproduced at least once (n= 555), 46.5% reproduced without being seen helping previously. Among all the females that reproduced at least once in our population (n= 590), 74.4% did not help before breeding. However, this sample includes many immigrant females first captured as breeders, which is likely to underestimate prior helping behaviour. Among resident females (n = 283), the proportion of individuals who bred without first helping dropped to 49.1%, a value close to that of males.

Males showed greater flexibility in reproductive trajectories than females considering the whole female data set. They were more likely to follow dynamic transitions between breeding and helping roles than females (Fig. 1 and 3; Table S1). Specifically, 13.6% of males followed a helper-breeder-helper path (vs. only 1.1% of females; simulated values ≈ 7.2% for both sexes; Fig. 3); and 8.5% followed a breeder– helper–breeder path (vs. 2.4% of females; simulated values ≈ 5.4%; Fig. 3). Males were also slightly more likely to transition from helper to breeder than females (17.4% vs. 14% of females; simulated values ≈ 15.7%; Fig. 3) and from breeder to helper (1.4% vs. 0.7% of females; simulated values ≈ 1%; Fig. 3). However, when focusing only on resident females, the results indicated higher flexibility in those females. Specifically, 47.4% followed the helper–breeder trajectory (compared to 14% in all females; Fig. S1; Table S1).

Although individuals were grouped into four combined reproductive strategies, some showed repeated status changes across time. On average, males switched between helper and breeder roles 0.99± 1.71 (mean ± SD) times across seasons, while females did so nearly 4 time less (0.23 ± 0.56). One male switched roles at least 15 times across seven breeding seasons. The most flexible female was resident in our population and switched five times in eight seasons, while the most flexible non-resident female changed roles four times across three seasons. We also recorded within-season switching, which occurred on average 0.59 ± 0.96 times in males and ca. 5 times less frequently in females (0.10 ± 0.39). The most flexible individuals changed roles up to five times within a single season (one male), four times (one non-resident female), and three times (one resident breeding female).

## Discussion

In this study, we used 10 years of data to describe breeding and helping dynamics in sociable weavers. We found that 66% of males and 40% of females help at least once in their life, that helping was common but not needed to obtain a breeding status (around 50% in resident birds helped before breeding). We uncovered that males helped for longer in their lives than females (approximatively twice as long), and more frequently, females being more likely to have only one reproductive status (i.e., breeder). Only half of both males and females were able to breed at least once, whereas ∼17% were never observed helping or breeding. Finally, we found remarkable flexibility in reproductive strategies in both sexes, but also marked sex-differences, with males being four times more likely than females to switch between helping and breeding roles between-seasons and about six times more likely within-seasons — a frequency of within-season switches that, to our knowledge, has not been documented in cooperative species. The sex differences in helping patterns and in the post-reproductive helping found here align with the fact that most females disperse to breed while males are philopatric, and is expected in other species with sex-biased dispersal and spatially close breeding sites

### Sexual differences in helping dynamics

Males were relatively more likely to help at least once than expected by chance than females. This is consistent with philopatric males having large opportunities to help kin [19,22], but our results reveal that these opportunities are used throughout life, long after males become reproductively active. Females, the dispersing sex, helped relatively less overall than expected by chance, probably because of reduced helping opportunities (as they have less relatives and social associations in the colonies they move into), and higher dispersal and reproductive costs [24, 37]. By contrast, resident females showed clear evidence of helping before breeding. Because we defined resident females as individuals that eventually bred, we did not capture trajectories of those that acted exclusively as helpers, and thus the frequency of resident females that helped at least once may be slightly underestimated. We also note that here we cannot distinguish between helping decisions and opportunity to help (e.g., if the subsequent breeding attempts of potential helpers’ parents did not reach the nestling stage). However, there is no reason to believe that the potential lack of opportunities would affect differently the results for males and females. Hence, these sex differences, and the helping by sociable weaver females, is notable given that the dispersing sex often helps little in cooperative breeders [13–14,25 but see 15], suggesting that extreme sociality and spatial breeding proximity promote cooperation.

### Transitions between breeding and helping and sex differences

Flexibility was marked in both sexes, but especially in males, switching between breeder and helper roles about four times more frequently than females. Although it was beyond the scope of this paper to examine the proximate causes of switching roles (e.g., failed breeding or poor breeding conditions), these patterns are consistent with males having greater opportunities and potential benefits from helping, even after breeding. Being philopatric and more closely related genetically to other males in the colony [27-28], males likely have access to more opportunities to help kin, as well as other long-term social partners, gaining indirect fitness benefits and potential reciprocal returns [15]. Recent evidence from superb starlings shows that individuals can help those who previously helped them, regardless of kinship [15]. Although starlings switch roles up to three times more often than weavers over their lifetime [15], such switches rarely occur within a single season [38]. By contrast, sociable weavers frequently reversed roles within a season, revealing unusual short-term plasticity. This flexibility may allow individuals, particularly males, to gain indirect fitness under fluctuating environmental conditions (e.g., rainfall; [39]) and variable food availability [29, 40-41]), which were shown to limit reproduction in this species [42]. Flexibility could enhance both direct fitness— through increased offspring production [43-45]— and indirect fitness by helping kin [46-47]. Indeed, studies in this population show that helpers increase brood survival [32] and improved the survival of young breeding females [3, 35].

Females also showed some flexibility, but markedly less than males. Although some females displayed role switching— one switched five times in eight seasons and another four times in a single season— most remained in a fixed breeder position. This likely reflects lower potential benefits of helping after dispersal (fewer related individuals around) and higher reproductive costs (e.g., egg laying). Indeed, females were three times more likely to never help than males. Altogether, these results suggest an evolutionary trade-off where dispersal costs combined with greater investment in reproduction and lower helping benefits, mediated by the females’ social environment, promote less flexibility in females in our population.

In conclusion, we reveal marked flexibility and sex differences in helping and reproductive strategies in sociable weavers, and an extremely long helping activity beyond the onset of independent breeding. Such behaviours are described mainly in mammals (linked to the evolution of menopause [48]). Here, we show that they also occur in a cooperative bird. These findings are consistent with sex-biased dispersal and social structure in this species and have implications for understanding post-reproductive helping across species. Further studies are needed to determine whether this flexibility is common and to quantify the associated fitness benefits and costs.

## Acknowledgments

We are very grateful to the numerous people involved in the long-term data collection over the years, especially for help with the breeding monitoring, annual captures and video analyses. The sexing and genotyping analyses were conducted at the CTM lab, CIBIO, and we thank Susana Lopes and all the staff that conduct the work. De Beers Mining Corporation provided access to Benfontein Nature Reserve and logistical assistance. This study was funded by the following grants: ERC-Consolidator 866489 (EU), FCT IF/01411/2014/CP1256/CT0007 and PTDC/BIA-EVF/5249/2014 (Portugal) to RC; ANR 15-CE32-0012-02 and 19-CE02-0014-01 (France) to CD. ACF was funded by University of Zurich Forschungskredit postdoc grant (K-74312-01-01 University of Zurich), Swiss Federal Commission for Scholarships and by ERC grant 850859 (awarded to Damien Farine). R.C. was further funded by FCT CEECIND/03451/2018). The Sociable Weaver project has received support from the and DST-NRF Centre of Excellence at the Fitzpatrick Institute of African Ornithology (University of Cape Town), the French OSU OREME, and the CNRS programme on Long-Term Studies in Ecology and Evolution (SEE-Life).

## Ethics Statement

All protocols followed the ASAB/ABS guidelines for the Use of Animals in Research (ASAB/ABS, 2018) and were approved by the Northern Cape Nature Conservation (permit FAUNA 1638/2015, 0825/2016, 0212/2017, 0684/2019 and 0059/2021) and the Ethics Committee of the University of Cape Town (permit 2014/V1/RC and 2018/V20/RC).

## SUPPLEMENTARY MATERIAL

**Table S1.**
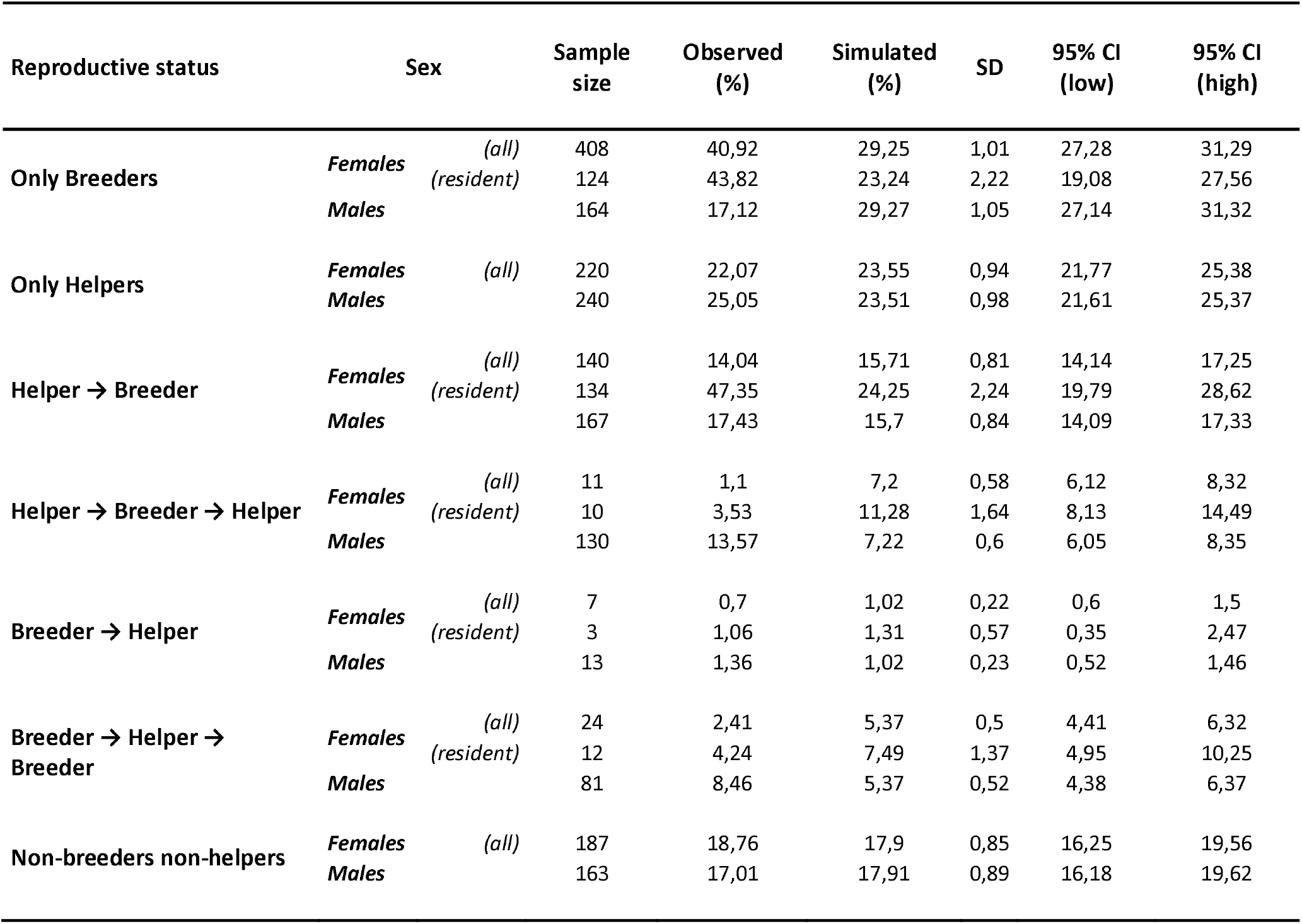
Simulated probabilities, observed percentages, and associated confidence intervals for each reproductive strategy (i.e., breeders, helpers, non-breeders-non-helpers, first breed and then help, first breed then help, and breed again, first help and then breed, and first help then breed, and help again) in the sociable weaver population. Observed percentages reflect cumulative data across the study period (from 2014/2015 to 2020/2021), representing the lifetime occurrence of each status. Simulated probabilities, along with standard deviations and variances, estimate the likelihood of individuals adopting each strategy across their lifetime.

| Reproductive status | Sex |  | Sample size | Observed (%) | Simulated (%) | SD | 95% CI (low) | 95% CI (high) |
| --- | --- | --- | --- | --- | --- | --- | --- | --- |
| Only Breeders | **Females** | (all) | 408 | 40,92 | 29,25 | 1,01 | 27,28 | 31,29 |
|  |  | (resident) | 124 | 43,82 | 23,24 | 2,22 | 19,08 | 27,56 |
|  | **Males** |  | 164 | 17,12 | 29,27 | 1,05 | 27,14 | 31,32 |
| Only Helpers | **Females** | (all) | 220 | 22,07 | 23,55 | 0,94 | 21,77 | 25,38 |
|  | **Males** |  | 240 | 25,05 | 23,51 | 0,98 | 21,61 | 25,37 |
| Helper → Breeder | **Females** | (all) | 140 | 14,04 | 15,71 | 0,81 | 14,14 | 17,25 |
|  |  | (resident) | 134 | 47,35 | 24,25 | 2,24 | 19,79 | 28,62 |
|  | **Males** |  | 167 | 17,43 | 15,7 | 0,84 | 14,09 | 17,33 |
| Helper → Breeder → Helper | **Females** | (all) | 11 | 1,1 | 7,2 | 0,58 | 6,12 | 8,32 |
|  |  | (resident) | 10 | 3,53 | 11,28 | 1,64 | 8,13 | 14,49 |
|  | **Males** |  | 130 | 13,57 | 7,22 | 0,6 | 6,05 | 8,35 |
| Breeder → Helper | **Females** | (all) | 7 | 0,7 | 1,02 | 0,22 | 0,6 | 1,5 |
|  |  | (resident) | 3 | 1,06 | 1,31 | 0,57 | 0,35 | 2,47 |
|  | **Males** |  | 13 | 1,36 | 1,02 | 0,23 | 0,52 | 1,46 |
| Breeder → Helper → Breeder | **Females** | (all) | 24 | 2,41 | 5,37 | 0,5 | 4,41 | 6,32 |
|  |  | (resident) | 12 | 4,24 | 7,49 | 1,37 | 4,95 | 10,25 |
|  | **Males** |  | 81 | 8,46 | 5,37 | 0,52 | 4,38 | 6,37 |
| Non-breeders non-helpers | **Females** | (all) | 187 | 18,76 | 17,9 | 0,85 | 16,25 | 19,56 |
|  | **Males** |  | 163 | 17,01 | 17,91 | 0,89 | 16,18 | 19,62 |

**Figure S1.**
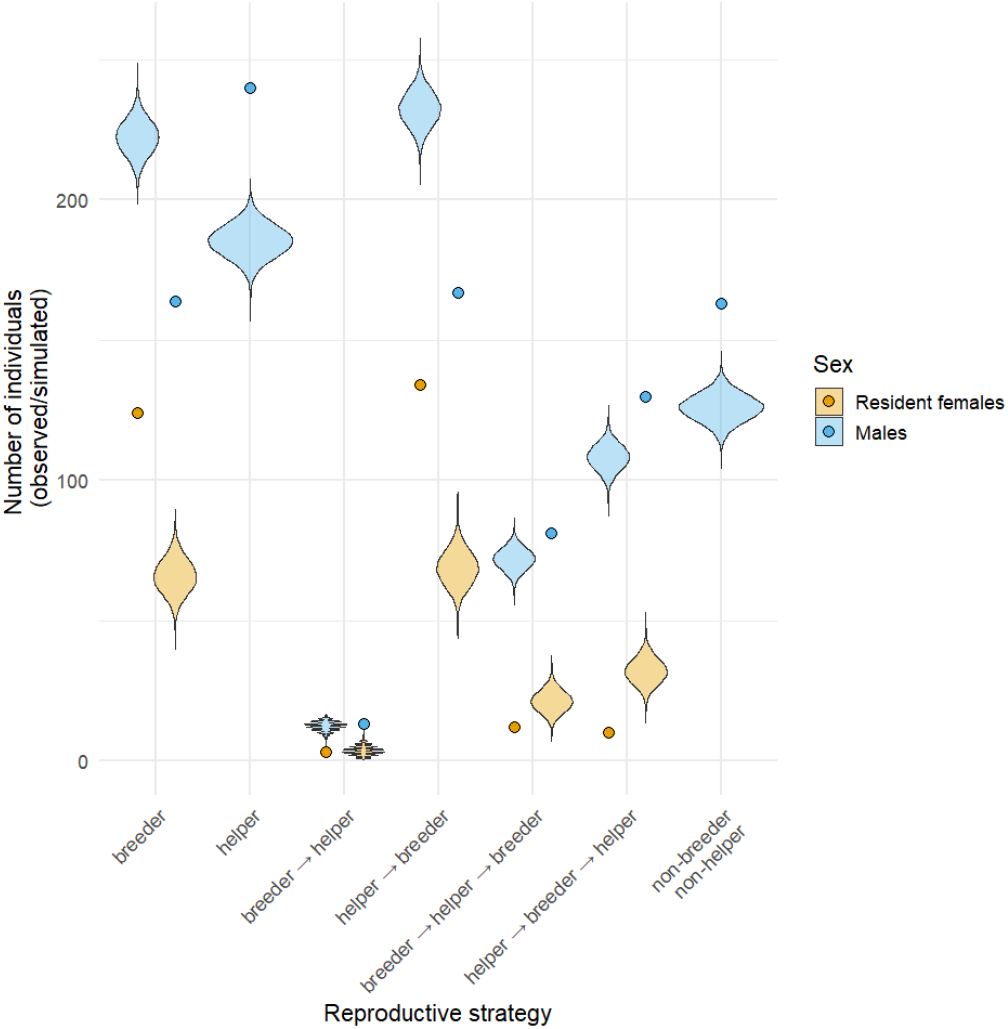
Observed and simulated numbers of individuals for each reproductive strategy in sociable weavers including only the resident females (i.e., those born and subsequently breeding in our study population) and all the males and. Violin plots show expected counts from 10,000 simulations under the null model. Points indicate the observed values. Colours designate sex (blue = males, orange = resident females). Note that the categories non-breeder-non-helper and helper only are not applicable for resident females (born and bred in the population).

## Supplementary Material & Methods

### Determination of Reproductive Strategies

Together with genetic parentage analysis, we analysed 6480 videos recorded during incubation and nestling provisioning to determine the breeder or helper status of 8656 visiting individuals across 1959 laying attempts between 2014 and 2023.

To establish the reproductive strategies (i.e., breeders, helpers and non-breeder-non-helper individuals) that males and females followed during each breeding attempt we used both breeding history, behavioural observations and genetic relatedness (see [1-2]). First, paternity was determined for breeders using a combination of: i) genetic markers (see [3-4]), ii) video analysis (see [1] for details), and iii) chambers’ reproductive history. Molecular estimates were robust and calculated in reference to genotypes from the entire population [4] (Van Dijk et al. 2015). We use COLONY Software v2.0 3.5 [5] (Jones and Wang 2010) to assign each chick the most likely mother and father. In addition, we also determined social fathers (i.e., male breeders) by using the behavioural data in cases where the genetics gave us two different fathers for any clutch (i.e., mostly due to errors, as only the 2.2% of young were extrapair in our population; [2] D’Amelio et al. 2024a). To identify helper individuals, we used video recordings from food provisioning. These videos provided a very effective way of obtaining the identity of the carers trough their colour ring combinations and measuring their contributions to nestling feeding and incubation [6] (Ferreira et al 2025). We considered as helper if an individual was seen visiting at least two times a given nest breeding event, in order to distinguish from prospecting individuals.

### Estimation of individuals’ age

Because of dispersal, exact age was not possible to assess for all the individuals. Age of the individuals that were not born in our colonies was estimated by attributing the average minimum age of first dispersal at their first capture, which is of 727 and 690 days for females and males, respectively [7] (Silva et al. 2025).

